# Sub-cellular Systems Drift Drives Mosaic Evolution of Mammalian Neurons

**DOI:** 10.64898/2026.03.27.714927

**Authors:** Jean Rosario, Junhyong Kim

## Abstract

Evolution of the mammalian brain has been described as mosaic evolution wherein natural selection for behavioral function promotes independent evolution of specific functional units despite developmental constraints that might govern overall change^1,2^. Evidence of mosaic evolution has been reported at the level of gene expression in individual structures^3^, cell type abundances^4^, as well as gene regulatory changes at the single cell level^5–7^. In particular, it has been hypothesized that brain evolution involves changes in circuit organization^6,8^. Circuit-level changes involve sub-cellular compartments that mediate synaptic activity, raising the question whether mosaic brain evolution might be found at the sub-cellular scale. Here, we examine the rate of evolutionary divergence between *Mus musculus* (C57BL/6) and *Rattus norvegicus* (Sprague–Dawley) for their dendritic transcriptome, which shapes the post-synaptic proteome through sub-cellular localization and local translation^9^. We address the problem of variable assessment of the dendritic transcriptome by micro-dissecting individual hippocampal pyramidal neurons to create matched single cell libraries of the soma and the dendrites from the same cell and apply a machine learning model to predict localization. Our results show that the dendritic transcriptome is significantly more divergent than the soma, but the core functional roles of the dendritically localized genes are conserved. Examining gene families for their localization suggests enrichment of family level conservation or localization. We propose that the observed divergence may arise from a combination of adaptive modulation and system drift under selection for core function. Our study suggests fine-grained mosaic evolutionary dynamics at the scale of synaptic function that mediates information processing and neural connectivity.

## Single cell matched sub-cellular sequencing and machine learning model uncovers high-confidence dendritically localized transcriptome in mouse and rat hippocampal neurons

Following our previous work dissecting matched dendrites^i^ and soma from the same single neuron^10^, we generated matched single sub-cellular transcriptome libraries from 16 individual cultured pyramidal hippocampal rat neurons (Sprague-Dawley) and re-quantified our previous C57BL6 mouse data to integrate the two species data (**Fig. 1a** and **1b**). On average, we detected 8,287 genes in mouse somas and 4,600 in rat somas, and 5,176 genes in mouse dendrites and 4,408 in rat dendrites. Read counts were normalized separately for each species using DESeq2’s size factor normalization, with cell identity and subcellular compartment included as batch factors^11^.

**Fig. 1:**
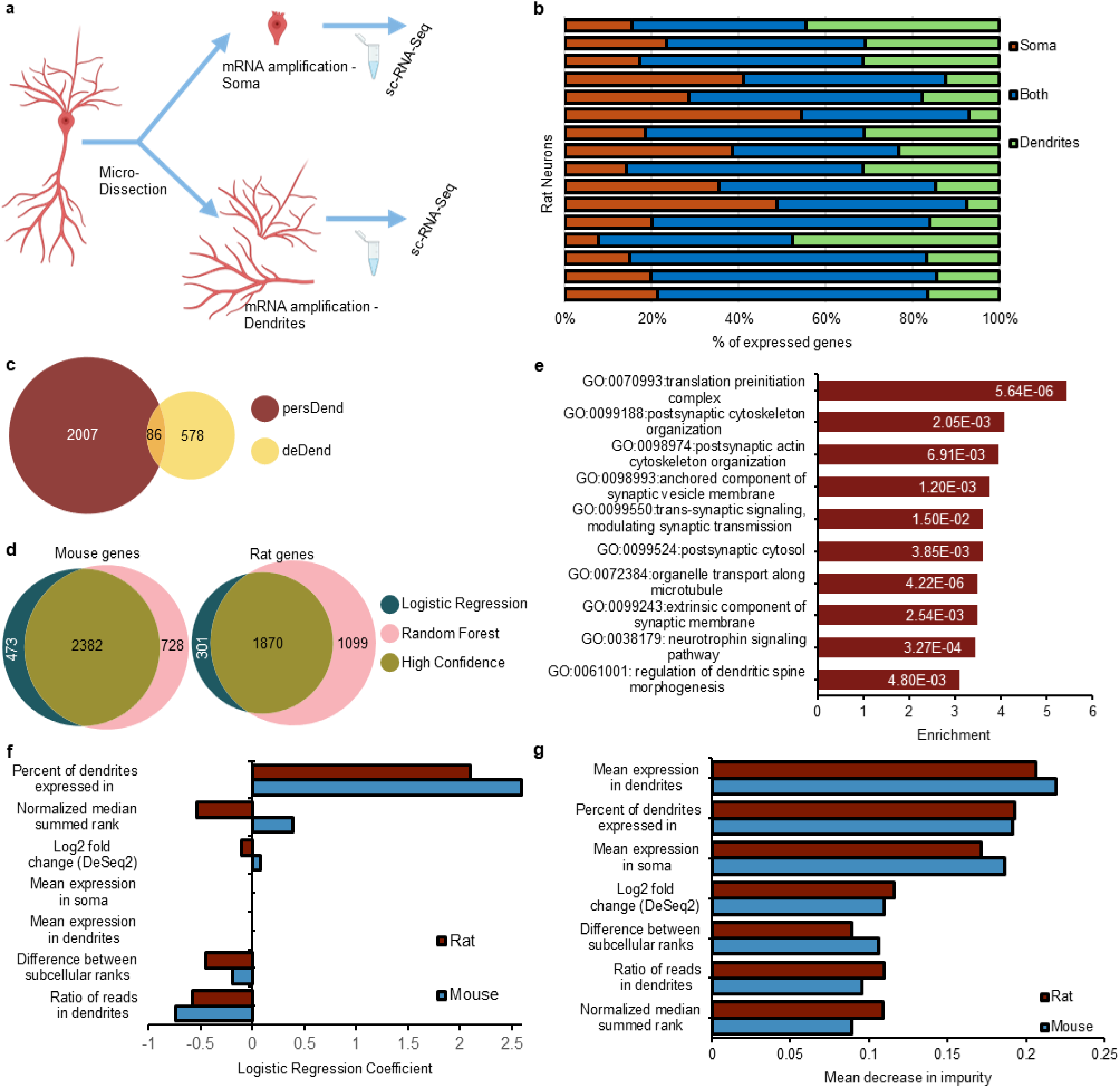
Subcellular sequencing of rat neurons and classification of localized genes. **a**, Soma and dendrites were separated from each rat neuron and collected utilizing a micromanipulator-controlled syringe with a glass micropipette, then individually amplified and sequenced. **b**, Overlap of genes expressed at greater than 10 reads between soma and dendrites for each individual neuron (each horizontal bar). **c**, Low overlap between persistently dendritic (persDend) and differentially expressed genes in dendrites (deDend). **d**, Overlap between the genes identified in the logistic model and the Random Forest in both mouse and rat. **e,** GO terms enriched in rat persDend genes. **f**,**g**, Informative expression statistics in the logistic regression (**f**) or Random Forest (**g**).

Following the analysis in previous mouse work using within-same cell differential expression analysis, we found 664 genes enriched in rat neuronal dendrites, which we call the deDend class^10^. We utilized Middleton et al’s persistently localized criteria to identify 2,093 transcripts that might not be dendritically enriched but are found to be consistently present in rat dendrites. We refer to this class as the persDend set (renamed from consDend in the original publication) (**Fig. 1c**). We also found minimal overlap between deDend and persDend, consistent with previous mouse results (**Fig. 1c**)^10^. Notably, GO terms associated with persDend genes are strongly enriched for dendritic and synaptic functions, indicating that persistent localization captures functionally relevant transcripts (**Fig. 1e**).

Assays of dendritic transcriptomes have high variability^10^. To overcome this, we applied logistic regression and Random Forest (RF) machine learning model to predict localization from expression features. Our logistic regression model identified 2,250 and 2,960 genes localized in rat and mouse neurons, respectively, whereas the random forest (RF) model identified 3,058 and 3,257 dendritically localized genes. Consistency of detection across dendrites emerged as the strongest positive predictor of localization, resulting in most persDend genes being classified as localized (**Fig. 1f, 1g,** and **Supplemental Fig. 1a**, **Supplemental table 1**). The gene set identified by the logistic regression are specifically enriched in dendritically related GO functional terms, supporting their functional relevance (**Supplemental Fig. 1b**).

The RF classifier identified 3,059 localized genes in rat and 3,258 in mouse (**Fig. 1d**, **Table 1**, and **Supplemental Fig. 1a**). While consistency of dendritic expression remained an important predictor, the RF classifier placed more importance on the mean expression in dendrites of each gene, as measured by the mean decrease in impurity (MDI) (**Fig. 1g**). This suggests that the RF classifier likely identified genes with high but variable dendritic expression, potentially reflecting context-dependent localization. Although genes unique to the RF model were not enriched for GO terms associated with distinct synaptic functions compared to the soma background, they showed enrichment for ribosomal and endoplasmic reticulum-related proteins, many of which are themselves localized, potentially reflecting roles in local translation and protein production (data not shown)^12,13^.

**Table 1.** Conserved and divergent expression of ortholog pairs.

| Expression gene set | Conserved | Mouse only expression | Rat only expression | Total orthologs | Jaccard similarity | Overlap coefficient |
| --- | --- | --- | --- | --- | --- | --- |
| Dendritically localized orthologous pairs sets |  |  |  |  |  |  |
| persDend | 757 | 780 | 1196 | 2,733 | 0.2770 | 0.4925 |
| deDend | 12 | 439 | 616 | 1,067 | 0.0112 | 0.0266 |
| Logistic Regression | 1261 | 1994 | 1024 | 4279 | 0.2947 | 0.5519 |
| Random Forest | 1338 | 2233 | 1890 | 5461 | 0.2450 | 0.4145 |
| High confidence | 960 | 1511 | 847 | 3318 | 0.2893 | 0.5313 |
| Expressed orthologous pair sets |  |  |  |  |  |  |
| Somatic fraction | 4955 | 5156 | 410 | 10521 | 0.4710 | 0.9236 |
| Hippocampus whole neurons | 7544 | 10002 | 957 | 18503 | 0.4077 | 0.8874 |
| Cortical whole neurons | 7492 | 5264 | 1150 | 13906 | 0.5388 | 0.8669 |

Both the logistic regression and RF classifiers emphasized consistency of dendritic detection as the strongest predictor, validating the need for our approach of single-neuron sub-cellular measurement. We defined a “high-confidence” set of localized genes as the intersection of the RF and logistic regression predictions within each species, resulting in 2,632 localized genes in mouse and 1,923 genes in rat (**Supplemental Fig. 1a**).

## Dendritic transcriptome shows rapid divergence compared to soma but maintains conserved functional ensembles

To dissect the evolution of dendritic localization across mouse and rats, we examined the pattern of localization of homologs between the two species. A notable percentage of genes in mouse and rat map to multiple homologous genes in the other species^14^. To address this ambiguity, we considered the localization status of all homologous pairs between genes expressed in rat and mouse neurons, rather than limiting to those that have a one-to-one relationship (**Fig. 2a**).

**Fig. 2.**
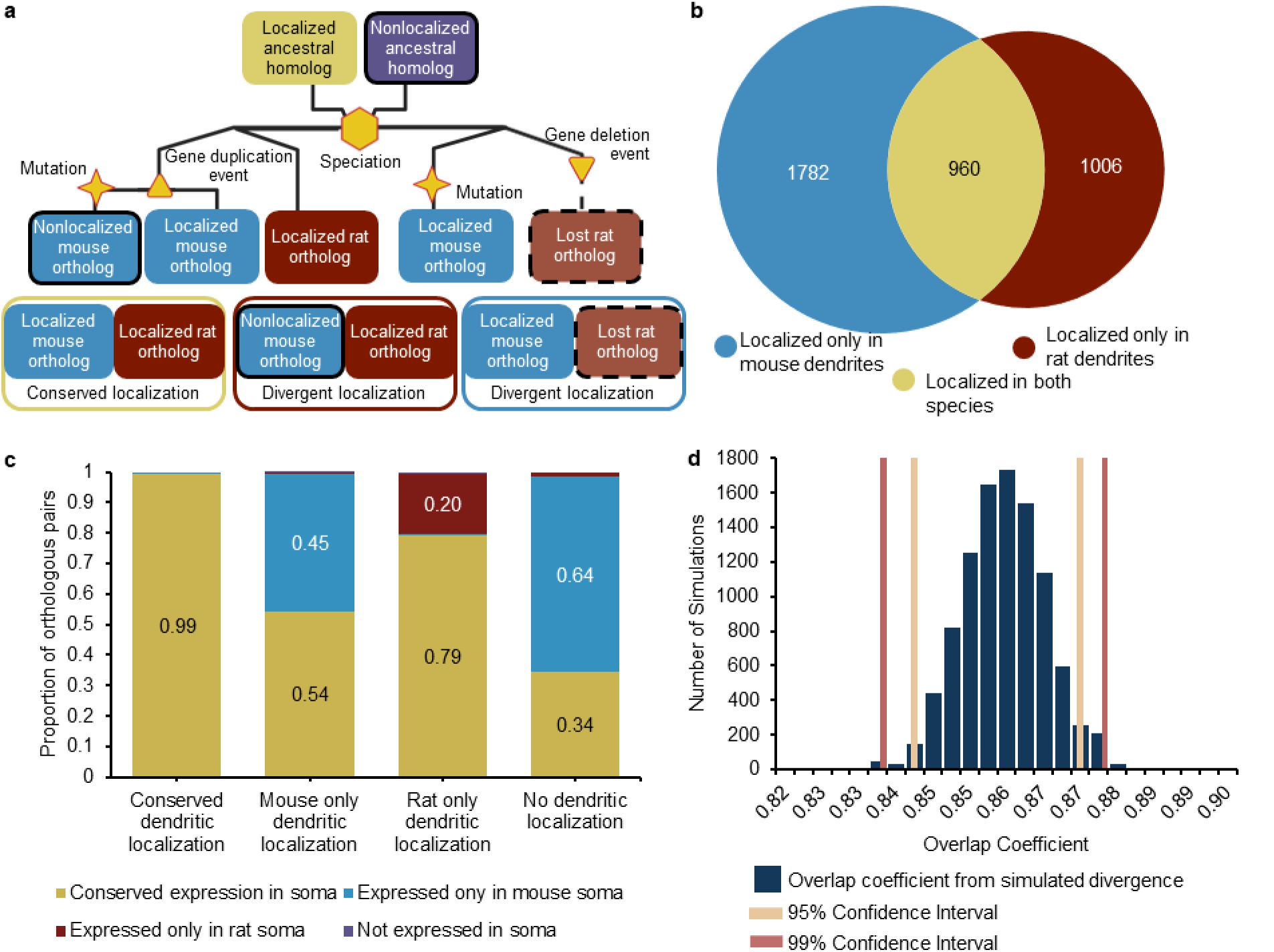
Divergent dendritic localization of orthologous gene pairs. **a**, Schema to assess conserved versus divergent dendritic localization in cases of one-to-many or many-to-many orthologs. **b**, Species overlap of the High confidence localized genes when considering genes or orthologous pairs. **c**, Expression of dendritically localized genes in their respective soma (expression classified as having 1 read in at least 50% of somatic samples). Divergent in dendritic expression is not likely due to divergence in somatic or whole cell expression. **d**, Similarity distributions were computed by using the genes expressed in at least half the cells of each dataset and randomly selecting sets of genes of similar size to the total High confidence gene set for localized genes. Dashed bounds and red bounds mark the 95% and 99% confidence intervals respectively.

Although this broader approach inflates the number of localized orthologous pairs and overlooks phylogenetic distance among orthologs, it enables more consistent cross-species comparisons and provides a fuller view of the evolutionary dynamics shaping the dendritic transcriptome^15,16^.

Comparing dendritic localization between mouse and rat neurons revealed limited conservation, with only 960 of 3,328 localized orthologous pairs dendritically localized in both species, an overlap coefficient of 0.407 (960 conserved pairs / 1,807 localized in rat, the smaller set), and Jaccard index of 0.2893 (960 / 3,328 localized orthologous pairs in at least one species) (**Fig. 2b** and **Table 3**). This limited overlap is not unique to the High-confidence set: persDend and deDend genes show overlap coefficients of 0.4925 and 0.0273, and Jaccard indevrd of 0.2770 and 0.0112, respectively (**Table 1**). The extremely low deDend overlap is largely driven by rat genes with low dendritic expression that are mostly absent from the soma, potentially due to incomplete RNA capture during microdissection (**Supplemental Fig. 2a-c**). Nevertheless, we observed low conservation of dendritic localization across all dendritically localized gene sets.

We compared dendritic divergence to baseline expression divergence in the corresponding somatic fractions from the same cell (**Fig. 2c**). Genes were considered expressed if they had at least one normalized read in 50% of somatic samples for each species, and ortholog comparisons were performed as described above. We observed far greater conservation in the somatic transcriptome, with an overlap coefficient of 0.9236 (Jaccard index 0.4710) (**Table 1**). While smaller, the rat somatic gene set (5,365 genes) is nearly fully encompassed by the mouse gene set (10,111 genes), indicating stable gene expression at the single-cell level despite incomplete genome annotations in the rat genome (**Table 1**). Re-analysis of single-cell hippocampal and cortical neurons from Dueck et al. (2015) further confirmed substantially higher conservation at the whole-cell level compared to dendrites, with overlap coefficients of 0.8874 and 0.8669, and Jaccard indices of 0.4077 and 0.5388, respectively) (**Table 1**)^17^.

To assess whether the observed dendritic overlap was lower than expected by chance, we ran 10,000 random sampling simulations matched to the size and composition of the High-confidence sets and calculated 99% confidence intervals for expected overlaps. Our simulation indicates that the dendritic transcriptome diverges more rapidly than can be explained by the whole-neuron and somatic expression differences (**Fig. 2d**). The loss of subcellular localization but conserved expression in whole-cell suggests that evolutionary changes are occurring at the level of mRNA cytoplasmic regulation and transport, consistent with subfunctionalization of orthologous genes across species with neofunctionalization at low but detectable rate^18,19^.

The observed divergence of dendritic transcriptome might imply evolutionary divergence in synaptic function. Comparison of mouse and rat electrophysiological properties have reported species-specific differences^20–23^. The observed differences in dendritic transcriptome might underlie this modulation of neuronal function. Nonetheless, most core synaptic properties remain highly conserved, as evidenced by functional neuronal circuits composed of mixed neurons from rats and mice^24,25^. Here we show orthologous pairs with divergent dendritic localization show substantially greater conservation at the functional level than between orthologs (**Table 1** and **Supplemental table 2**).

To assess whether the ensemble of localized transcripts is conserved at the level of biological function, despite differences in the identity of the genes, we examined the overlap of mouse and rat dendritic transcriptome in terms of their GO annotations. Using mouse GO annotations for both species, the overlap coefficient of GO terms associated with divergently localized orthologous pairs is 0.6693 (Jaccard 0.3808), and an overlap coefficient of 0.7443 (Jaccard 0.4473) when only considering neuronally-associated GO terms. (**Fig. 3a**, **b**, and **Supplemental Table 2**). Reactome annotations show even higher conservation, with an overlap coefficient of 0.7434 and a Jaccard similarity of 0.4473 (**Supplemental Table 2**)^26,27^. Random sampling of genes with divergent expression in somatic fractions or whole-cell datasets confirmed that GO-term similarity for divergently localized genes exceeds expectations from expression divergence alone (**Supplemental Fig. 3a**). GO terms shared between divergently localized orthologous pairs tend to occur higher in the GO hierarchy, i.e., more general functions, and likely represent core synaptic and dendritic molecular functions compared to species-specific GO terms, which tend to be lower in the hierarchy, representing more narrow functions (**Fig. 3b**). Taken together, these observations suggest a model of system drift for core functions, in which orthologous genes show differential gene-level subcellular localization while preserving core neuronal functions across species^28–30^. In the meanwhile, some differentially localized genes likely represent species-specific modulation of synaptic function.

**Fig. 3.**
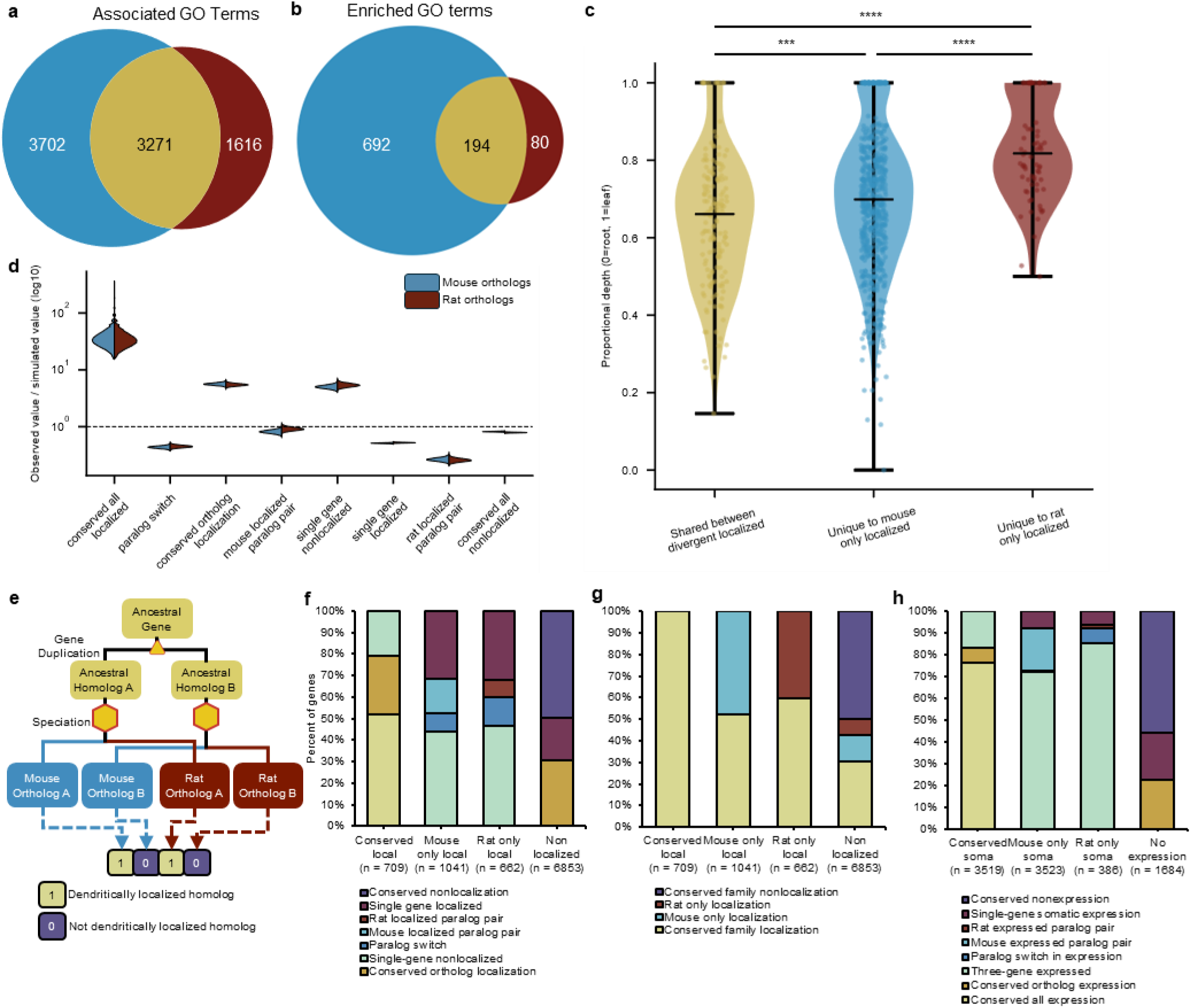
Conservation of core molecular functions. **a**,**b**, Overlap between all GO terms (**a**) associated with or (**b**) enriched in differentially localized orthologous pairs (0 overlap between pairs). **c**, Distribution of mean proportional depth for GO terms in **b**. Proportional depth is defined GO terms position in graph / total number of terms from root to terminal term. **d**, Distribution of observed quartet per gene counts / simulated quartet per gene evolutions from 10,000 random simulations of the evolution of 9,265 mouse and 8,996 rat gene families containing 4 members each. \*\*\**P* < 0.001, \*\*\*\**P* < 0.0001. **e**, Schematic of homolog quartets that are used for homolog family analysis. Each combination of homologs and paralogs were considered individually for each set of annotations. **f**, The evolutionary events that lead to specific localization classification. **g**, Evolutionary events that lead to specific expression in neuron somas. **h**, The broad family level localization due to observed patterns of localization evolutions in homologs.

## Paralog localization is enriched among divergently localized genes

Conservation of core function, despite gene-level divergence, could arise through paralog switching, that is the evolution of species-specific subcellular usage of paralogous genes, or paralog retention, where some homologs are not present in dendrites but the others are, preserving overall gene family expression as individual gene’s sub-cellular localization diverge^18,28,29,31,32^. To test these possibilities, we examined the localization patterns of the paralogs of each localized gene. Using Ensembl annotations, we identified paralogs in both species, incorporating relationships traced back to the Opisthokont divergence. To avoid complicated paralogous relationships we restricted our analysis to paralog-ortholog homologous quartets composed of four genes^14^.

In **Fig. 3e**, we coded each quartet of genes by their orthology/paralogy and localization status as a vector of four values. Here, position 1 and 3 are orthologs and 2 and 4 are also orthologs.

Position 1 is paralogous to position 2 and position 3 is paralogous to position 4. The vector is given a value of 1 (localized) or 0 (not localized), in respective species (**Fig. 3e** and **Table 2**). This binary quartet framework allows us to characterize the subcellular localization of the quartets by their binary patterns. For example, patterns 1101 or 0111 denote cases of paralog retention or loss, while 1001 or 0110 denote paralog switching.

**Table 2.** Categorization of binary vectors of homologous gene quartets.

| Mouse paralogs |  | Rat paralogs |  | Mouse gene counts | Mouse gene percent | Rat gene counts | Rat gene percent | Ortholog localization status | Quartet Localization Status | Gene family localization status |
| --- | --- | --- | --- | --- | --- | --- | --- | --- | --- | --- |
| 1 | 1 | 1 | 1 | 369 | 4.02% | 367 | 4.08% | Conserved localization | Conserved localization | Conserved family localization |
| 1 | 0 | 1 | 0 | 193 | 2.10% | 193 | 2.15% | Conserved ortholog local. | Conserved ortholog local. | Conserved family localization |
| 0 | 1 | 0 | 1 | 2009 | 21.90% | 2011 | 22.35% | Conserved ortholog local. | Conserved ortholog local. | Conserved family localization |
| 0 | 1 | 1 | 1 | 308 | 3.36% | 539 | 5.99% | Single gene nonlocalized | Single gene nonlocalized | Conserved family localization |
| 1 | 0 | 1 | 1 | 50 | 0.54% | 101 | 1.12% | Single gene nonlocalized | Single gene nonlocalized | Conserved family localization |
| 1 | 1 | 0 | 1 | 455 | 4.96% | 289 | 3.21% | Single gene nonlocalized | Single gene nonlocalized | Conserved family localization |
| 1 | 1 | 1 | 0 | 97 | 1.06% | 47 | 0.52% | Single gene nonlocalized | Single gene nonlocalized | Conserved family localization |
| 1 | 1 | 0 | 0 | 167 | 1.82% | 52 | 0.58% | Mouse only localized | Mouse only localized | Mouse family localization |
| 0 | 0 | 1 | 1 | 54 | 0.59% | 184 | 2.05% | Rat only localized | Rat only localized | Rat family localization |
| 1 | 0 | 0 | 1 | 89 | 0.97% | 89 | 0.99% | Paralog switch | Paralog switch | Conserved family localization |
| 0 | 1 | 1 | 0 | 89 | 0.97% | 92 | 1.02% | Paralog switch | Paralog switch | Conserved family localization |
| 1 | 0 | 0 | 0 | 330 | 3.60% | 208 | 2.31% | Single gene localized | Single gene localized | Mouse family localization |
| 0 | 1 | 0 | 0 | 841 | 9.17% | 499 | 5.55% | Single gene localized | Single gene localized | Mouse family localization |
| 0 | 0 | 1 | 0 | 211 | 2.30% | 381 | 4.24% | Single gene localized | Single gene localized | Rat family localization |
| 0 | 0 | 0 | 1 | 508 | 5.54% | 789 | 8.77% | Single gene localized | Single gene localized | Rat family localization |
| 0 | 0 | 0 | 0 | 3405 | 37.11% | 3155 | 35.07% | Conserved nonlocalized | Conserved nonlocalized | Conserved nonlocalized |
| Totals |  |  |  | 9175 | 100.00% | 8996 | 100.00% |  |  |  |

In over half of orthologous gene pairs with divergent dendritic localization, the gene that lacked dendritic localization retained a paralog that was dendritically localized in the same species. Of the quartets exhibiting divergent ortholog localization, approximately 45% reflected divergence driven by a single gene paralog losing or gaining dendritic localization, whereas about 29% arose from a single gain of dendritic localization in one gene (e.g the pattern 1000). In contrast, paralog switching appears to be relatively rare, making up only 10% of the cases of divergent dendritic localization (**Fig. 3f** and **Table 2**).

To test whether this apparent conservation of paralog localization exceeds expectations under random gain or loss of localization, we simulated localization evolution across ∼9,000 four-member homolog quartets, running 10,000 replicates to generate expected localization patterns under a fixed evolutionary rate of gain or loss of localization in dendrites (**Fig. 3d**). The ancestral localization frequency for each family was estimated by grouping homologs according to their ancestral lineage, as inferred by annotated homologous relationships. Using the simulated distributions of binary quartet configurations, we computed 99% confidence intervals to identify quartet states enriched or depleted relative to random expectation (**Supplemental Table 3**).

The simulation results revealed a clear deviation from random gain/loss. Quartet states preserving gene-family localization, where orthologs are localized (1010,1111, 0101) or only one quartet member is nonlocalized (e.g. 1011), were strongly enriched compared to all other categories. In contrast, paralog switching (1001 or 0110) was shown to be depleted in our data set. While patterns with localization only in rat were also rare (e.g. 0011), patterns with mouse only localization occurred more frequently than expected (**Fig. 3d**). Our data suggests that selection maintains dendritic function at the level of gene families, with paralogous genes providing compensatory capacity as individual members gain or lose localization. Performing the same homologous quartet analysis in the somatic transcriptome reveals far less single-gene expression and generally less functional differences in terms of gene family expression in the somatic compartment (**Fig. 3g**). The dendritic transcriptome remains more divergent than the somatic and whole neuron transcriptome, at both the level of orthologous pairs and gene-family.

Duplicated paralogs may be retained by undergoing subspecialization via changes in subcellular localization at either the mRNA or protein level^33–35^. Quartet states where three out of the four members are localized may represent partial or transitional events consistent with ongoing subfunctionalization, where paralogs have diverged in one species but not yet in the other^36,36–38^. Paralog switching requires both gain and loss of localization and the three-out-of-four transitional state might be part of the switching process. The genes participating in this paralog switching or partial switching quartets are enriched for GO terms related to synaptic and dendritic function suggesting that this process preferentially affects genes with dendritic importance (**Supplementary Table 5**). Divergence of localization at the gene level, coupled with conserved function, hints that paralogous genes may compensate for lost localization, a form of evolutionary system drift in dendritic transcriptomes.

## Discussion and Conclusion

In this study we generated a matched sub-cellular dendrite-soma single-cell RNA-seq dataset for rat hippocampal neurons and integrated the mouse data from Middleton et al (2019). We generated a comprehensive high confidence set of dendritically localized genes consisting of 2,632 genes in mouse and 1,923 in rat. Our data is one of the first robust catalogs of dendritic localization at subcellular resolution for cross-species comparison and evolutionary analysis.

Out of a total of 3,328 localized orthologous gene pairs in our High Confidence gene sets, only 960 pairs displayed conserved dendritic localization, an overlap coefficient of 0.5313 and Jaccard index of 0.2893 (**Fig. 2b** and **Table 1**). Compared with expression conservation in somatic compartments, overlap coefficient of 0.9236 and Jaccard index of 0.4710, or whole single-cell neuron datasets (overlap = 0.8874 and Jaccard = 0.4077) from Dueck et al. 2015, dendritic localization shows disproportionately high divergence (**Table 1**). Despite this gene-level turnover, we see much greater conservation in molecular and pathway-level annotations in the dendrites. Orthologous pairs with divergent localization still share significantly large overlapping GO and Reactome terms. Shared GO terms preferentially map to more basal functional annotations, suggesting conservation of core function, while species-specific GO terms preferentially map to more specialized functions, suggesting evolutionary modulation of synaptic function. (**Fig. 3c** and **Supplemental table 2**).

To probe how function persists despite localization divergence, we constructed paralog–ortholog “quartets,” containing the focal gene, a paralog, an ortholog, and a paralog of the ortholog. (**Fig. 3e**)^29,30,39^. We found that over half of divergent ortholog pairs retain a localized paralog in the opposite species, meaning that dendritic localization is preserved at the gene-family level even as individual orthologs diverge. Among quartets with divergently localized orthologs, ∼45% showed only one nonlocalized gene (e.g. 1011), ∼29% reflected a single gene being localized (e.g. 0010), and only ∼10% matched classical paralog switching (**Fig. 3f** and **3g**). Simulations modeling neutral localization drift demonstrate enrichment for quartet states with conserved family-level localization and depletion of species-specific or paralog-switching events. These patterns of paralog localization suggest that the preservation of dendritic localization at the gene-family level arises not through strict conservation of individual orthologs, compensatory shifts among related paralogs, with molecular function being distributed among localized paralogs or condensed on the individually localized gene.

It has long been proposed that extant paralogs persist through subfunctionalization, as one paralog is lost to genetic drift if both remain functionally redundant^19,31^. However, more recent studies reveal that homologous genes can retain overlapping molecular functions, enabling the underlying genotype to drift while maintaining stable phenotypes at both cellular and organismal levels^18,28^. One example of paralog maintenance without subfunctionalization is co-evolution of paralogs whose combined expression is under selection. a so-called compensatory drift model.

Compensatory drift, or system drift, could explain the cases where the localization of only one paralog in one species diverges, while the other species retains localization in both paralogs. (**Fig. 3d**, **3f**, and **3g**).

Studies have reported variation in molecular and electrophysiological properties of neurons, which could underpin functional differences in brain function^20–23^. However, studies of neuronal input–output relationships have revealed pronounced degeneracy in neuronal circuits, in which diverse molecular and cellular configurations can support similar functional behavior^40–43^. This degeneracy is thought to arise from compensatory molecular changes and activity-dependent plasticity, enabling neurons to buffer perturbations in individual components, including core synaptic elements such as ion channels that regulate electrical signaling^43–47^. We propose that when viewed in an evolutionary context, the degeneracy observed in neuronal circuits can be viewed as system drift, in which conserved neuronal functions are maintained despite substantial turnover in the underlying molecular and cellular components.

Our results support a model in which dendritic mRNA localization evolves through compensatory system drift, in which selection acts primarily on core dendritic function rather than on the precise identity of individual localized genes. Under this model, individual genes can gain or lose dendritic localization with limited fitness consequences, provided that functionally overlapping genes remain present in the dendritic compartment. Paralog redundancy and shared molecular functions therefore create the opportunity for compensatory shifts in localization, allowing extensive gene-level turnover while preserving dendritic molecular capacity.

## Methods

### Neuron culture and subcellular collection

Hippocampal neurons were collected from embryonic day 18 (E18) Sprague-Dawley rats and cultured as described by ^48^. Briefly, neurons were plated at 75,000 cells/mL in Neurobasal medium supplemented with B-27, on 12-mm round German Spiegelglas coverslips coated with either poly-L-lysine or poly-D-lysine. To match the timeline used by Middleton et al. (2019), rat neurons were collected after 15 days in culture, although 14 days are sufficient for extensive neurite maturation and expression of appropriate marker proteins ^10,49^. Cell culture density was reduced from 100,000 to 75,000 cells/mL to facilitate individual cell collection; no changes in cell morphology or maturation were observed ^50^.

Subcellular fractions were collected as follows: the soma was first separated from the dendrites of a neuron using a glass micropipette mounted on a micromanipulator under 10× magnification and immediately expelled into a collection tube. The dendrites from the same neuron were then collected by sweeping a second micropipette around the area previously containing the soma, taking care to avoid the neurites of adjacent neurons, and transferred into a separate tube. Each collection tube contained first-strand synthesis buffer and RNase inhibitor and was kept at 4°C to minimize RNA degradation.

A total of 28 rat neurons were collected across multiple days and animals; however, only 16 neurons passed quality control metrics in both their somatic and dendritic fractions and were used for downstream analyses.

### Subcellular RNA-amplification and sequencing

Poly-adenylated RNA from all subcellular fractions was selected and amplified using three rounds of aRNA in vitro transcription in ^51^. For quality control and quantification, each sample was spiked with ERCC control RNA, diluted to 1:4,000,000 final working concentration. Strand-specific sequencing libraries were prepared from the amplified material using the Illumina TruSeq Stranded kit, following the manufacturer’s instructions, except that the initial poly-A selection step was omitted since aRNA amplification already excludes non-polyadenylated RNA.Pooled libraries were sequenced across three independent runs on an Illumina NextSeq instrument using a 75 bp paired-end kit, achieving an average depth of 25 million reads per sample. Adapter and poly-A trimming were performed using an in-house processing pipeline, and reads were aligned to the rat genome (rn6) with STAR ^52^. From the STAR mapping, uniquely mapped reads were used for feature quantification, as gene expression counts, using VERSE (described below) ^53^.

### Gene expression counts and subcellular localization analysis

We combined the Ensembl genes annotations (downloaded from Ensembl Genome Browser, Dec. 2015) and UCSC genes annotations (downloaded from UCSC, Dec. 2015) to quantify reads with VERSE using options “-s 1 -z 3 --nonemptyModified” ^14,53,54^. Gene expression analysis was carried out separately for both rodent species, with a gene being considered “expressed” in the neurons of a particular species if it was present at a least one normalized read in half the collected samples for that species (32 total soma-dendrite pairs from a single neuron, 16 for mouse and 16 for rat). These neuronally “expressed” genes were then used for differential expression analysis and to identify non dendritically localized genes that are nevertheless expressed in rodent neurons.

DESeq2 was used both for library size normalization, via the size-factor method, and for identifying differentially expressed genes between soma and dendrite fractions using a paired experimental design. This design incorporated cell identity as a batch factor for normalization across the somatic and dendritic samples of the same cell ^11^. Significantly differentially expressed genes were identified using a false discovery rate (FDR)-corrected p-value < 0.05, defining the deDend gene set in each species. Each species’ persDend gene set was defined as genes with at least one size-factor–normalized read in ≥90% of the collected dendritic samples within that species (15 of 16 samples) ^10^.

### Identification of dendritically localized genes through machine learning

To perform the categorization of localized genes through supervised machine learning, we first compiled a list of 4 localization studies for each species, For mouse we used data from ^55–58^. For rat, we used ^58–61^. Genes present in at least two studies within a species were designated as “true positives” for localization, while genes which were absent from all localization studies but detected with at least one read in one sample were considered “true negatives.” This resulted in 287 and 182 localized (“true positive”) genes for mouse and rat, respectively, and 532 and 368 nonlocalized (“true negative”) genes for each species.

The known dataset for each species was split such that 75% of both localized and nonlocalized genes were used to train the models, and the remaining 25% (71 mouse genes and 40 rat genes) were reserved for testing. Logistic regression and random forest (RF) classifiers were implemented using Scikit-learn ^62^. Logistic regression was run with default parameters and limited to 1,000 iterations; models failing to converge within this limit were considered unsuccessful. Several expression-derived statistical features were tested, and the seven metrics shown in **Supplemental table 1** yielded the highest classification accuracy, correctly identifying approximately 60% of true positives and 90% of true negatives. The RF classifier was trained using 100 decision trees and the same seven features, training, and test sets as the logistic regression model. Model comparisons were performed both at the species level and across finer localization categories, **Supplemental fig. 1a**.

### Building orthologous pairs for localization comparison

As described above and highlighted in **Fig. 2a**, determining orthologous relationships between localized genes for evolutionary analysis was non-trivial and required detailed comparison of annotated orthologs between rat and mouse. While we primarily relied on orthology annotations from the Ensembl database, we supplemented these relationships with in-house gene symbol matching, identifying orthologs by pairing identical orphan gene symbols lacking Ensembl IDs across species^14^.

### GO ontology analysis of dendritically localized genes

Functional categorization of localized genes was approximated by gene ontology (GO) enrichment analysis, as calculated by the GOrilla webserver or the PANTHER ^63,64^ The enriched set of GO terms was selected by an FDR-adjusted p-value ≤0.05.

For mouse, the background set of genes was identified as the genes present in at least 90% of neurons, defined by the summed reads of the dendritic and somatic pair samples, while in rat the background set of genes were present in only 80% of neurons. The lower threshold in rat was selected to account for reduced genome annotation quality in rat, which otherwise resulted in substantially fewer detectable genes. The 80% threshold in rat yielded a comparable annotation recovery (∼29%) to that obtained in mouse at the 90% threshold (**Supplemental fig. 3**).

While analyzing conserved or divergently dendritically localized gene sets, neuronally-associated GO terms were selected as those containing the one of the following key words in the GO terms description: neuron, neuronal, axon, dendrite, dendritic, projection, synapse, synaptic, nerve, nervous, myelin, growth cone, RNA localization, or establishment of localization. While this list is overly conservative, it seeks to exclude very general terms, such as translation and ion channels, that might be overly broad and artificially increase the expected GO overlap within localized and nonlocalized genes within a species and divergently localized genes between species.

### Simulation of expected overlap for the dendritic transcriptome and associated GO terms

To examine whether the observed dendritic divergence could arise simply from partitioning the neuronal transcriptome into compartments, we simulated randomly taking a subset the neurite transcriptome and computed the Jaccard coefficients (Intersection / Union) and Overlap coefficients (Intersection / Smallest set, i.e., the maximum possible overlap between two sets) for randomly drawn gene sets. Each simulated set was of comparable size to the union of the dendritic transcriptomes of both species (3,318 genes).

For each simulation, we randomly drew 3,318 ± 300 pairs from the 10,521 pairs expressed in the soma of at least one species. We then used the previously recorded expression status of each pair (conserved expression in both species, expression only in mouse neurons, or expression only in rat neurons) to compute the Jaccard and overlap scores. The resulting distributions were generated from 10,000 iterations to establish the 95% and 99% confidence intervals shown in **Fig. 2d**.

Similarly, to estimate the expected overlap of gene ontology (GO) terms associated with neuronal functions among divergently localized and expressed genes, we mapped the neuronal GO terms associated with the unique genes in each of the randomized drawings above (i.e., the symmetric difference between the two sets) across iterations. The distribution of this simulation is shown in **Supplemental fig. 4a**.

### GO proportional depth analysis

Gene Ontology (GO) term depth analysis was performed to characterize the hierarchical specificity of GO terms associated with dendritically localized gene sets and genes undergoing paralog switching. GO term annotations were retrieved using the full GO ontology graph parsed from the go-basic.obo file (Gene Ontology Consortium), and hierarchical depth metrics were computed by traversing the GO directed acyclic graph (DAG) for each term^65,66^. For each GO term, all possible path lengths to the root node and to terminal leaf nodes were enumerated via depth-first search, yielding distributions of root distance and leaf distance. These distances were used to compute a proportional depth, defined as the ratio of root distance to total path length (root-to-leaf) for each path through the term, where values near 0 indicate proximity to the root (general function) and values near 1 indicate proximity to terminal leaves (specific function).

From these path distributions, minimum, maximum, mean, and median values were computed for each metric per term. Analyses were performed separately for each GO namespace (Biological Process, Molecular Function, Cellular Component) as well as across all namespaces combined. Group differences in depth distributions were assessed using the Kruskal-Wallis test across all groups, followed by pairwise Mann-Whitney U tests. All analyses were implemented in Python using the goatools library for GO DAG parsing, with scipy for statistical testing^67,68^.

### Estimating evolution rates of dendritic mRNA localization and ancestral

To simulate the evolution of localization under neutral genetic drift, we first estimated the ancestral frequency of localized genes. Because it is unlikely that neuronal dendritic and synaptic functions remained completely unchanged throughout early evolution, we assumed a lower ancestral localization rate than observed in rat and mouse. To estimate ancestral localization rate, we clustered all related paralogs in each species and computed the mean proportion of localized gene per cluster as a basis for our ancestor state localization frequency, which was approximately 120/4422 based on mouse paralog annotations and 124/4292 based on rat paralog annotations.

The most likely rate of evolution (*p*) of dendritic localization per species was estimated by maximizing the following likelihood function:

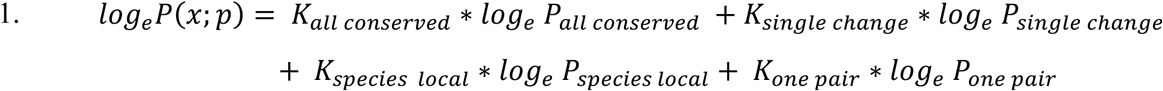

The identify of the equation above are as follows:

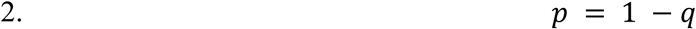

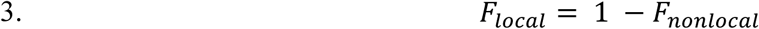

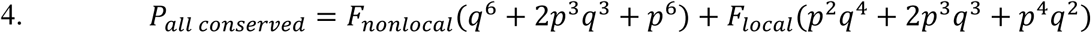

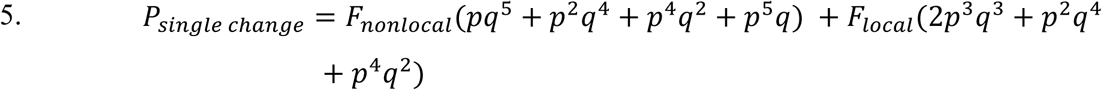

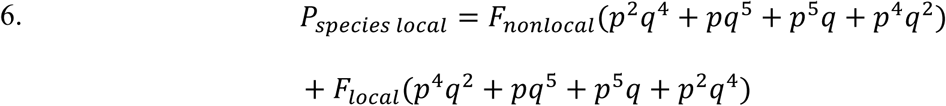

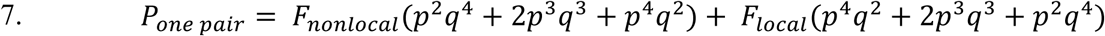

where 𝐹_𝑛𝑜𝑛𝑙𝑜𝑐𝑎𝑙_ and 𝐹_𝑙𝑜𝑐𝑎𝑙_ are the ancestral frequencies of nonlocalized and localized genes, respectively, and *K* represents the sum of homologous quartet vector counts observed in the data:

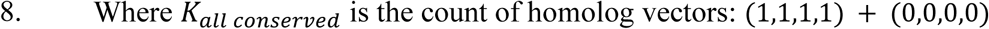

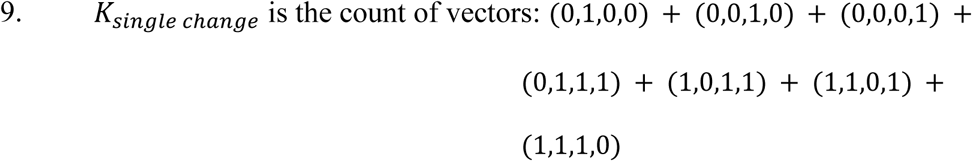

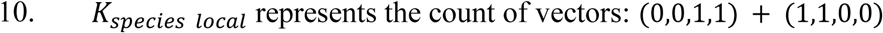

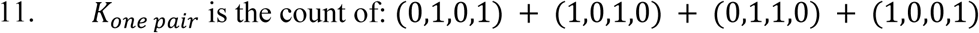

From the above identities we can arrive at:

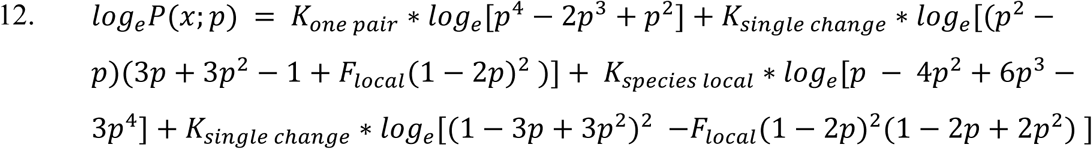

Taking the first derivative of equation [12] yields the log maximum-likelihood estimate of *p*, the rate of dendritic localization evolution. From our observed data, we estimate:

𝑝_𝑚𝑜𝑢𝑠𝑒_ ≈ 0.174149, 𝑝_𝑟𝑎𝑡_ ≈ 0.177963

The small difference between the two estimates reflects differences in the number of annotated paralogs per species, which lead to slightly different ancestral cluster counts and homologous quartet totals.

### Random simulations of genetic drift in the evolution of dendritically localized genes

We used the estimates of ancestral localization frequency and evolutionary rate of novel localization to simulate the evolutionary history of 9,265 independent homologous gene quartets over 10,000 times. While all 9,265 homologous gene quartets are not fully independent in the observed data, this simplification allow us to compute the simulations more easily and fully randomizes the evolution of localization in the quartets. Briefly, the probability of the ancestral gene being localized was set equal to the estimated ancestral localization frequency. Following gene duplication events, the probability of a change in localization status (localized → nonlocalized, nonlocalized → localized) was set according to the species-specific evolutionary rates (𝑝_𝑚𝑜𝑢𝑠𝑒_ or 𝑝_𝑟𝑎𝑡_ above). SSpeciation between rat and mouse then generates the full homologous quartet, and the change in localization status is again applied according to the estimated evolutionary rate. From these simulations, we calculated the 99% confidence bounds for each binarized homologous quartet, which were then used to identify significant deviations from drift in the observed data. Simulation results are summarized in **Table 2**.

## Footnotes

i Dendrites and axons are not morphologically distinguished for hippocampal pyramidal neurons at this stage, however vast majority of the neurites are dendrites and we use the term dendrites as approximation.

