## Supplemental Figures 1 - 4, Supplemental Tables 1 -3 for "Sub-cellular Systems Drift Drives Mosaic Evolution of Mammalian Neurons"

#### Supplemental Fig. 1

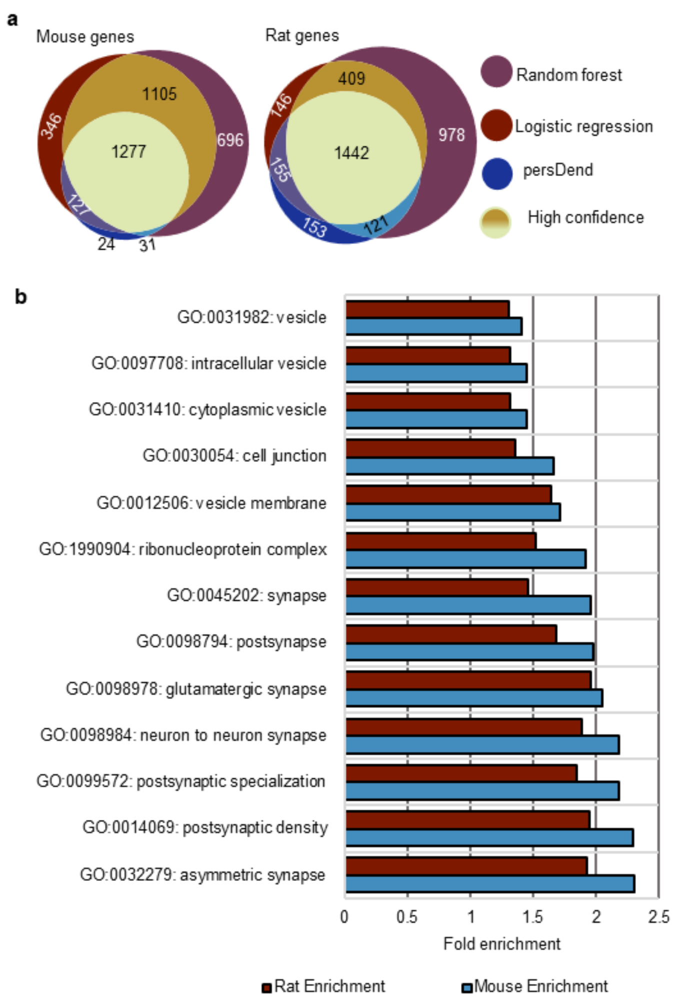


**a**, Overlap in different models of localization across species, the overlap of the Random forest and Logistic regression comprised the High confidence gene set. **b**, Neuronal function related GO terms found in genes identified by logistic regression.

**Supplemental Fig. 2 deDend genes low expression in dendrites**

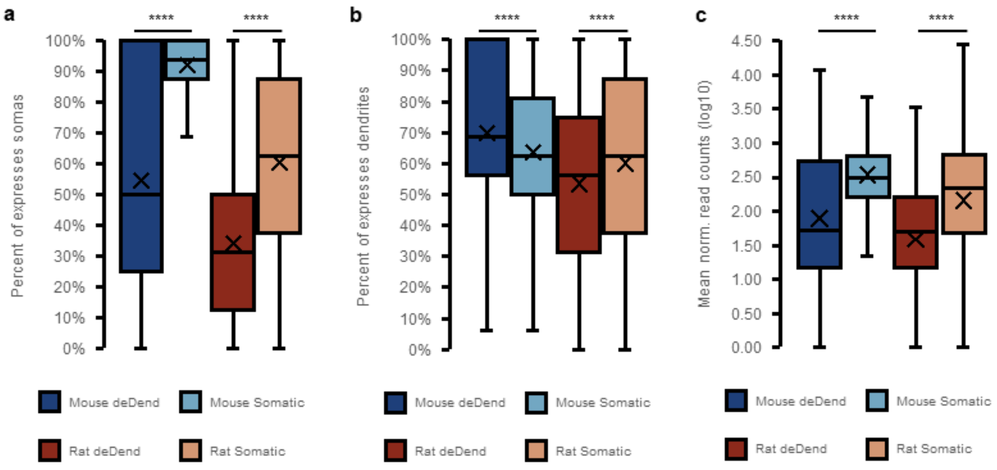


**a-b**, Individual sub compartment expression of deDend, differentially expressed in dendrites, compared to not deDend genes (termed somatic here). Genes were considered expressed in each individual sub compartment if at least one normalized read was detected. **c**, Mean normalized read counts for genes in the dendrites and somatic fraction. deDend genes tended to have lower expression in overall compared to somatic genes, likely leading to genes expressed highly in both compartments being misclassified as somatic.

**Supplemental Fig. 3 GO term overlap of divergently localized orthologous pairs**

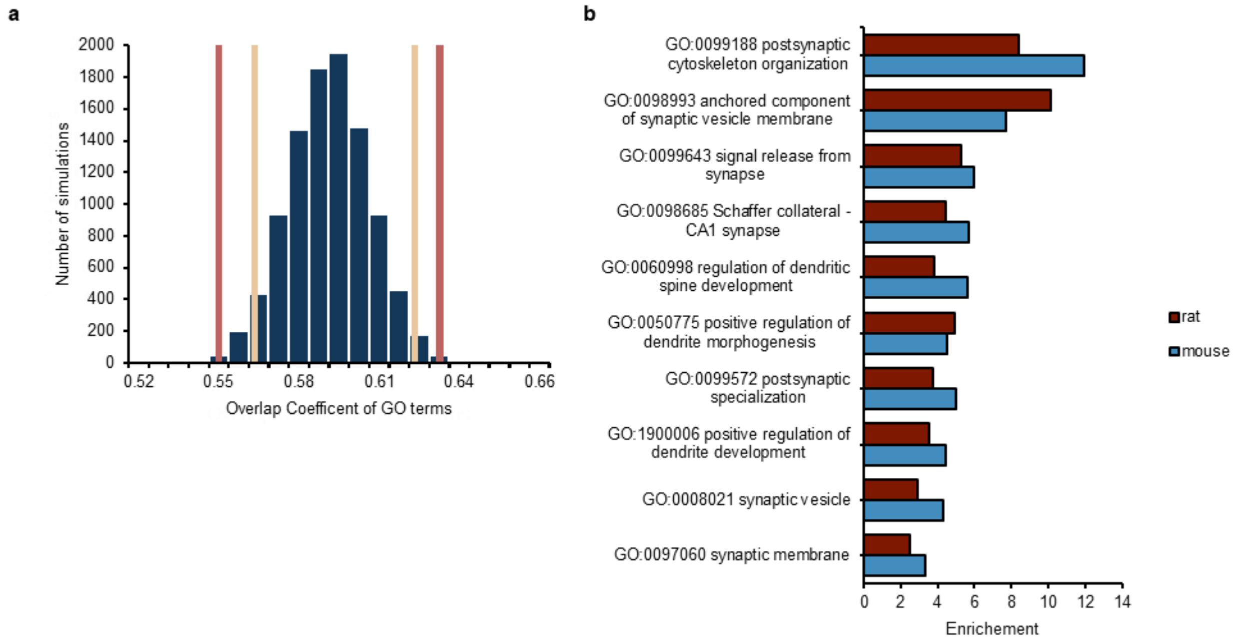


**a**, Similarity distributions for the overlap of GO terms associated with divergently expressed genes. Orthologous pairs were chosen at random from the divergently expressed genes in soma to match the size of the divergently localized ortholog pairs in **Fig. 2b**. **b**, Examples and enrichment of neuron-related GO terms shared by and unique to differentially localized orthologous pairs (from **Fig. 3b**)

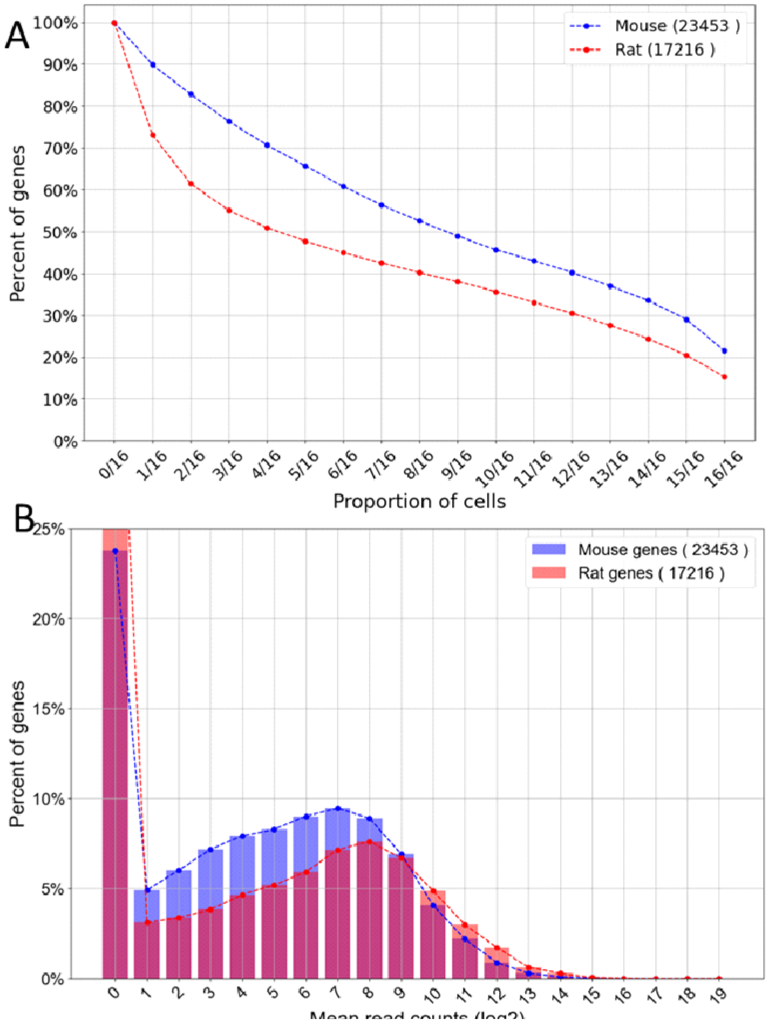


**Supplemental fig. 4: Percentage of genes recovered after size factor normalization.**

**a**, Gene dropout rate between mouse (blue) and rat (red). Genes were counted as being expressed in a cell if it contains at least 1 normalized read in either compartment. **b**, The read per gene distribution between mouse (blue) and (red), rat tends to have less expressed genes with larger number of reads per gene, suggesting missing rat gene annotations that lead lower number of overall rat genes.

### Supplemental tables

**Supplemental table 1 Expression statistics used for both logistic regression and random forest machine learning models to identify localized genes.**

| Expression statistics | Description |
|---|---|
| mean somatic expression | mean gene normalized expression counts in somatic compartments |
| mean dendritic expression | mean gene normalized expression counts in dendritic compartments |
| mean summed expression | mean of the summed gene normalized expression counts in the two compartments |
| mean dendrite to soma ratio | mean value of normalized dendritic reads / normalized somatic reads, calculated per dendrite soma pair |
| percent dendritic expression | The percentage of dendrites the gene is found at least 1 normalized read |
| percent somatic expression | The percentage of somas the gene is found at least 1 normalized read |
| mean dendrite all ratio | mean value of the proportion of reads in the paired neuron that were in the dendritic sample, calculated per dendrite soma pair |
| mean dendritic to soma difference | mean value of the difference between normalized dendritic reads and normalized somatic reads, calculated per dendrite soma pair |
| median dendritic to soma difference | median value of the difference between normalized dendritic reads and normalized somatic reads, calculated per dendrite soma pair |
| median dendrite all ratio | median value of the proportion of reads in the paired neuron that were in the dendritic sample, calculated per dendrite soma pair |
| median somatic expression | median gene normalized expression count in somatic compartments |
| median dendritic expression | median gene normalized expression counts in dendritic compartments |
| median summed expression | median of the summed gene normalized expression count in the two compartments |
| median dendrite to soma ratio | median value of normalized dendritic reads / normalized somatic reads, calculated per dendrite soma pair |
| count dend count soma ratio | percent dendritic expression / percent somatic expression |
| log2fold | Fold enrichment as computed by a paired design in DeSeq2 (described above) |
| median dendritic rank | Median expression rank in each dendrite. Reverse order ranking was used to assign ranks, with ties having the same rank |
| median somatic rank | Median expression rank in each soma. Reverse order ranking was used to assign ranks, with ties having the same rank |
| median rank differences | Median rank differences between dendrites and soma. Rank differences were computed per paired subcellular sample of the same neuron |
| mean dendritic rank | Mean expression rank in each dendrite. Reverse order ranking was used to assign ranks, with ties having the same rank |
| mean somatic rank | Mean expression rank in each soma. Reverse order ranking was used to assign ranks, with ties having the same rank |
| mean rank differences | Mean rank differences between dendrites and soma. Rank differences were computed per paired subcellular sample of the same neuron |
| differences median rank | Difference between median dendritic rank and median somatic rank |

**Supplemental table 2 Overlap of functional terms associated with divergent localized**

| Functional annotations | Shared | Mouse only | Rat only | Total orthologs | Jaccard similarity | Overlap coefficient |
|---|---|---|---|---|---|---|
| GO terms – Mouse | 3271 | 3702 | 1616 | 8589 | 0.3808 | 0.6693 |
| GO terms – Rat | 3048 | 3403 | 1699 | 8150 | 0.3740 | 0.6421 |
| Reactome terms – Mouse | 1476 | 797 | 195 | 2468 | 0.5981 | 0.8833 |
| Reactome terms – Rat | 1474 | 721 | 248 | 2443 | 0.6034 | 0.8560 |

**Supplemental Table 3: Rodent homolog quartet condensing and total numbers.**

| Category | Mouse condensed homologous quartets | Mouse total homologous quartets | Rat condensed homologous quartets | Rat total homologous quartets |
|---|---|---|---|---|
| Dendritically localized | 1750 | 23234 | 1346 | 16202 |
| Dendritically nonlocalized | 7515 | 1506908 | 7650 | 258020 |
| Somatically expressed | 7042 | 430806 | 3703 | 684293 |
| Somatically non-expressed | 2070 | 19445433 | 5490 | 19201,528 |
| Whole neuron expressed | 10846 | 3101467 | 5627 | 716436 |
| Whole neuron non-expressed | 845 | 17849215 | 6373 | 20305500 |
